# Assessing the impact of transcriptomics data analysis pipelines on downstream functional enrichment results

**DOI:** 10.1101/2023.09.13.557538

**Authors:** Victor Paton, Attila Gabor, Ricardo Omar Ramirez Flores, Pau Badia-i-Mompel, Jovan Tanevski, Martin Garrido-Rodriguez, Julio Saez-Rodriguez

## Abstract

Transcriptomics, and in particular RNA-Seq, has become a widely used approach to assess the molecular state of biological systems. To facilitate its analysis, many tools have been developed for different steps, such as filtering lowly expressed genes, normalisation, differential expression, and enrichment. While numerous studies have examined the impact of method choices on differential expression results, little attention has been paid to their effects on further downstream functional analysis using enrichment of gene sets, such as pathways, which typically provides the basis for interpretation and follow-up experiments. To address this gap, we introduce FLOP (FunctionaL Omics Processing), a comprehensive nextflow-based workflow that combines various methods for preprocessing and downstream enrichment analysis, allowing users to perform end-to-end analyses of count level transcriptomic data. We illustrate FLOP capabilities on diverse datasets comprising samples from end-stage heart failure patients and cancer cell lines in both basal and drug-perturbed states. We found that the correlation between gene set enrichment analysis results can vary significantly for alternative pipelines. Additionally, we observed that not filtering the data had the highest impact on the correlation between pipelines in the gene set space, especially in settings with limited statistical power. Overall, our results underscore the impact of carefully evaluating the consequences of the choice of preprocessing methods on downstream enrichment analyses. We envision FLOP as a valuable tool to measure the robustness of functional analyses, ultimately leading to more reliable and conclusive biological findings.

**Graphical abstract:** 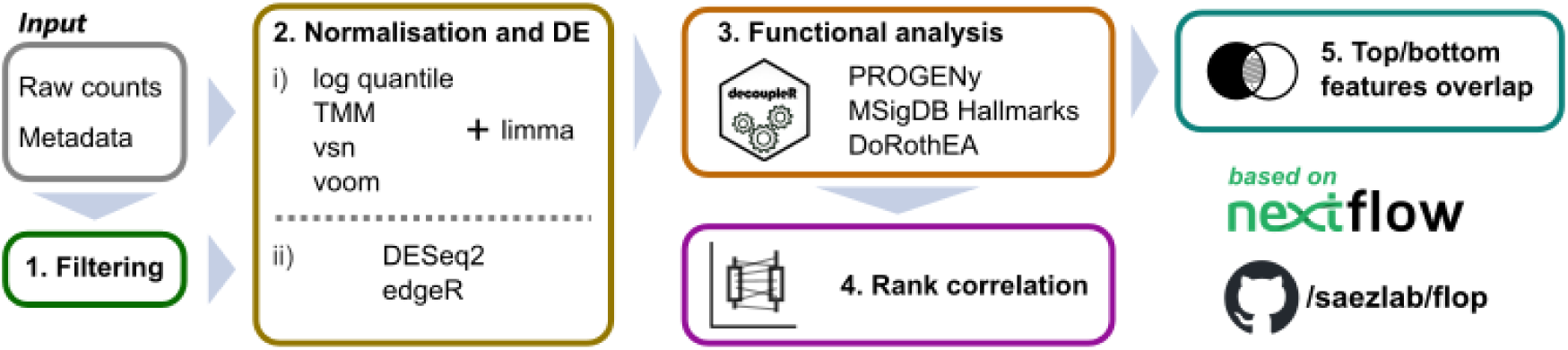

## 1. Introduction

Over the past years, advancements in sequencing technologies have allowed their democratisation across the scientific community [1]. In particular, RNA-seq offers the possibility to profile entire transcriptomes with nearly complete coverage in a cost-effective manner, and has become the reference technique to analyse gene expression states both at bulk and single-cell levels [2]. In the typical workflow to analyse bulk RNA-seq data, sequences are aligned to a reference genome or transcriptome, and the overlapping sequences within regions of interest (such as genes) are counted or estimated, [3–5]. This results in a count matrix, which is commonly filtered to exclude lowly expressed genes. The counts are then normalised to minimise the influence of technical effects and to make samples comparable. Usually, differential expression (DE) analysis is conducted afterwards, followed by a functional enrichment analysis based on gene sets of processes of interest such as pathways [6,7].

When filtering genes with low expression levels, researchers assume that variance in the measurements for these genes is mostly due to technical noise [8]. However, it is also possible that genes in the low expression range still encode valuable biological information, and hence there are different approaches to deal with them. During the filtering step in transcriptomics data analysis, researchers make a choice between coverage and accuracy, yet it remains unknown how this choice impacts downstream analyses.

The normalisation step is necessary to eliminate systematic biases that can impact the comparison of gene expression values between samples [9,10]. These biases can originate from a variety of sources, such as different library sizes, varying RNA content during different cell cycle phases and the well-studied mean-variance relationship that affects most sequencing technologies. There are multiple methods that have been adapted or developed to carry out the normalisation of RNA-Seq count data. Log quantile normalisation [11,12] and variance stabilising normalisation (vsn) [13] were initially designed for microarray data but later adapted for RNA-Seq data. On the other hand, methods like the trimmed mean of M-values (TMM) or voom were designed to be used specifically with RNA-seq data [9,10]. All DE methods require a previous normalisation step, and it has been shown that the choice of a specific normalisation method can significantly affect the resulting DE analysis results [14,15].

The goal of DE analysis is to compare the normalised expression measurements between samples. Also in this step, there are multiple available methods, and three of them are commonly included in most pipelines: Limma [16], edgeR [17] and DESeq2 [18]. Limma requires a previous normalisation step and models gene expression as a linear function of variables that encode the experimental design. edgeR and DESeq2 directly model count-level data using parametric models that assume an underlying negative binomial distribution of the data. Such models are then used to perform the comparison between samples [9,19].

DE analyses output results for thousands of genes, making direct biological interpretation challenging.[20]. To reduce dimensionality and enhance interpretability of DE results, genes are usually grouped into functional categories. This step is commonly referred to as functional enrichment analysis or gene set enrichment analysis. These functional categories, also known as gene sets, represent biological processes such as signalling cascades, metabolic pathways or genes that respond to specific chemical or genetic perturbations [21].

Early functional analysis methods primarily focused on over-representation analysis (ORA), which are typically conducted after thresholding DE results to determine which genes significantly change their expression levels. One of the first and most widely used ORA tools was DAVID [22], which provided users both gene set collections and a web application to carry out the analyses. Unlike ORA, Gene Set Enrichment Analysis (GSEA) [23] evaluates the entire gene set instead of individual genes, and does not require a threshold to determine deregulated genes. In recent years, consensus methods have also emerged as a way to combine the results from multiple functional analysis strategies, leading to the development of tools such as Piano [24] or decoupleR [25].

The diverse array of alternative approaches for conducting various stages in the downstream analysis of RNA-seq data poses a considerable obstacle in establishing standardised pipelines. To overcome this challenge, benchmark studies (see Supplementary Table 1) were carried out to facilitate the selection of appropriate methods [14,15,26–33]. These benchmark studies employ different strategies to assess method performance, with many utilising qRT-PCR or simulated data to define a reliable reference for comparison.

Although these benchmarks have provided valuable insights, they focus on comparing individual methods for specific steps and do not investigate the cumulative effects of applying alternative methods in sequence. Secondly, the simulated data used in some benchmarks may inadequately capture the complexities and behaviours observed in real transcriptomic data. Lastly, most benchmark studies overlook the influence of method selection on functional analysis outcomes. Consequently, there is a need for consistent standards that help researchers understand how various methods influence data processing and interpretation in transcriptomic analyses [20].

To evaluate the impact of preprocessing methods choice on downstream functional results, we developed FLOP (FunctionaL Omics Processing), a nextflow-based [34] workflow that analyses transcriptomic data using multiple combinations of filtering, normalisation and DE methods, and which evaluates their agreement after functional analysis. In its default version, FLOP uses transcription factors, signalling pathways, and transcriptional hallmarks as functional categories. In the absence of a ground truth that would enable us to make a claim about the performance of different pipelines, we focused on providing an easy and reproducible way of assessing the impact of alternative methods in the robustness of downstream results.

## 2. Results

### 2.1 FLOP: A workflow to compare transcriptomic preprocessing pipelines after functional analysis

In this study, we evaluated alternative transcriptomics data analysis pipelines, conceived as combinations of methods that perform the entire processing from raw counts to gene set enrichment scores. Each method input and output are often not directly compatible, making it difficult to combine them for analysis. To address this, we developed FLOP (FunctionaL Omics Processing platform), a unified workflow that takes a count matrix and a metadata file as input and applies alternative processing pipelines to produce functional enrichment scores using different gene grouping categories (Figure 1).

**Figure 1.**
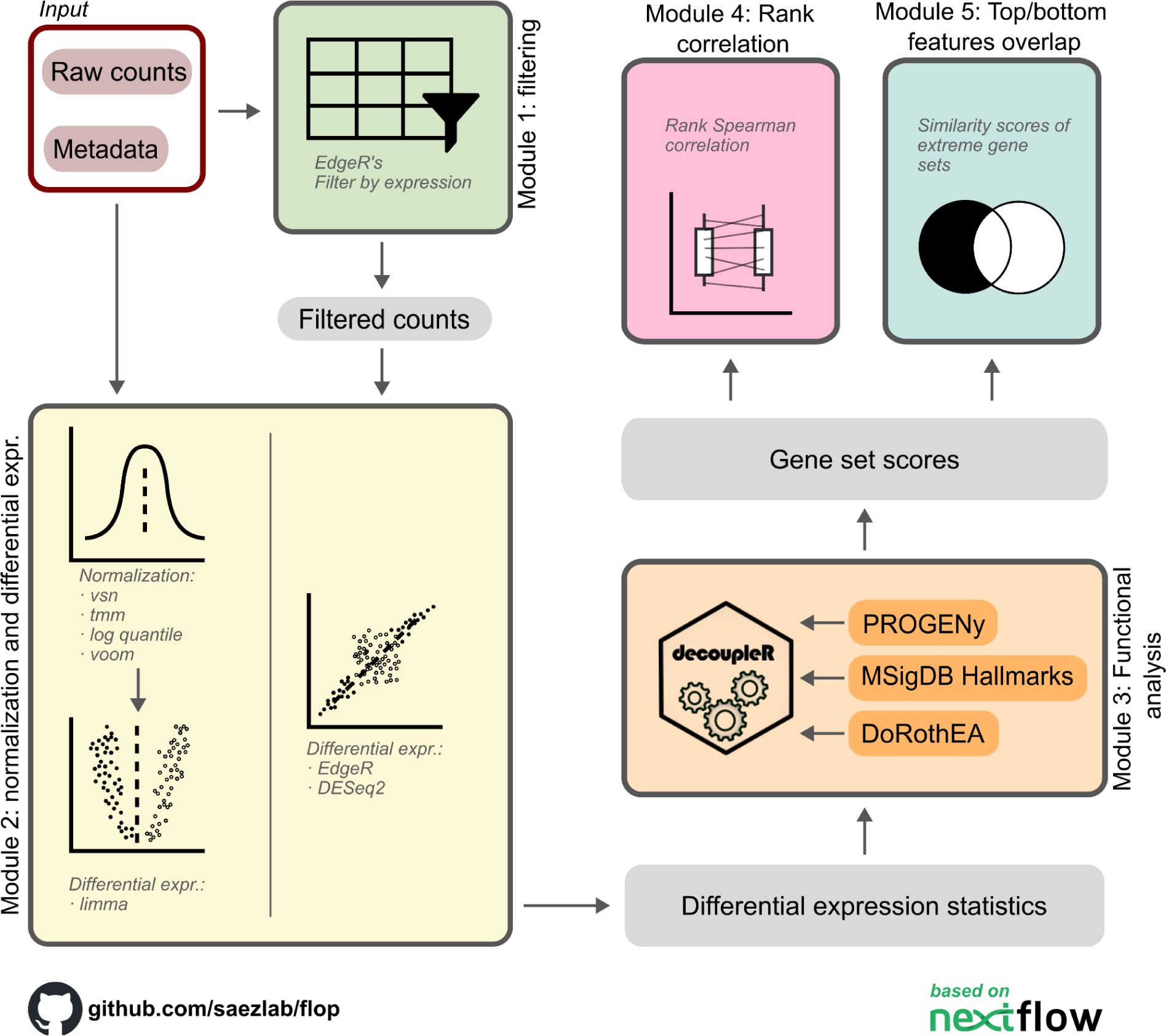
Schematic representation of FLOP methods and modules. From transcript counts and metadata, it uses several differential expression analysis frameworks, performs enrichment analysis and then evaluates the differences nt the reuutts iat woo different metrics.

To ensure that the analyses can be reproduced, extended, and scaled effectively, we developed FLOP using Nextflow [34,35]. We grouped the methods into three processing modules: (I) filtering, (II) normalisation and DE, and (III) functional analysis. We also created two evaluation modules (Figure 1). In the first module, the count matrix is either left intact or filtered using edgeR’s function *filterByExpr* with user defined parameters. The second module takes the filtered or non-filtered count matrix as input and applies multiple combinations of normalisation and differential expression (DE) methods. We coupled TMM, vsn, log quantile normalisation and voom normalisation with limma, in addition to edgeR and DESeq2, which employ raw counts as input. This results in a total of 6 combinations of normalisation and differential expression analysis tools that coupled to the filtering module creates 12 alternative pipelines.

We unify the output of each individual pipeline in a single table, which contains at least an effect size estimate (e.g. log fold change), a statistical estimate (e.g. t-values) and a significance estimate (e.g. adjusted p-value) of gene expression changes. As recommended by Zyla et al. [36], we use the statistical estimate as input for the functional analysis module. In the third module, we use the univariate linear model (ulm) method from the decoupleR package [25], along with three prior-knowledge sources storing different collections of gene sets: signalling pathway footprints (PROGENy, [37]), transcription factor targets (DoRothEA, [38]) and transcriptional hallmarks (MSigDB hallmarks, [39]). We chose the ulm method because it indicates at the same time the direction of the enrichment, positive or negative, and its significance in a single score, and also was observed to be stable and to outperform other methods in a previous benchmark [25].

Following the generation of enrichment scores for each pipeline, contrast, and functional category, we apply the evaluation modules. The "Rank correlation module" (Figure 1) assesses the consistency of different pipelines across the complete list of gene sets. To achieve this, the Spearman rank correlation is computed for pairs of pipelines, resulting in a single score that quantifies the similarity in rankings of gene sets within biological contrasts. In the fifth module, denoted as the "Top/bottom features overlap," we account for the possibility that some researchers may concentrate on the highly up-regulated and down-regulated gene sets for a given contrast. Consequently, this module examines the overlap between the top and bottom sections of the lists through the utilisation of a similarity index. The number of features to compare is a user-defined parameter.

To showcase the capabilities of FLOP, we applied it to datasets studying different biological contexts and with varying sample sizes. In the first case, we utilised datasets used for a recent meta-analysis, which focused on studying the transcriptomic profiles of patients with end-stage heart failure [40]. We then applied FLOP to analyse a large transcriptomic dataset that evaluated the basal transcriptomic profiles of various types of cancer cell lines [41]. Finally, we employed FLOP to analyse the recently published PANACEA DREAM challenge data, which explored the transcriptional response of 11 cell lines to 32 kinase inhibitors [42].

### 2.2 Application of FLOP to end-stage heart failure transcriptomic studies

We first applied FLOP to five transcriptomics datasets collected from the ReHeaT resource [43–47], a compendium of case-control bulk transcriptomics datasets of human heart failure profiling the left-ventricle of patients with dilated or ischemic cardiomyopathy and control myocardium from healthy donors.

To exemplify the different data modalities generated and compared during FLOP execution, we focused on a single study (Spurrel et al. [44]), a single contrast (heart failure versus control samples), and on the comparison of two specific pipelines: filtered counts analysed with DESEq2 versus unfiltered counts analysed with edgeR. As expected, the total number of genes and differentially expressed genes is higher in the pipeline on which the count matrix was not filtered. Specifically, when filtered, the total number of analysed genes is 16,117, compared to the 35,229 that compose the original matrix. Similarly, 638 genes are differentially expressed in the filtered data, while 1,151 are found to surpass the same cutoffs in the unfiltered data (adjusted P < 0.05 and absolute logFC > 1) (Figure 2A). If we compare the DE results in the common gene space, that is, genes that are scored in both pipelines, we observe a perfect correlation between them (Spearman correlation = 1) (Figure 2B).

**Figure 2.**
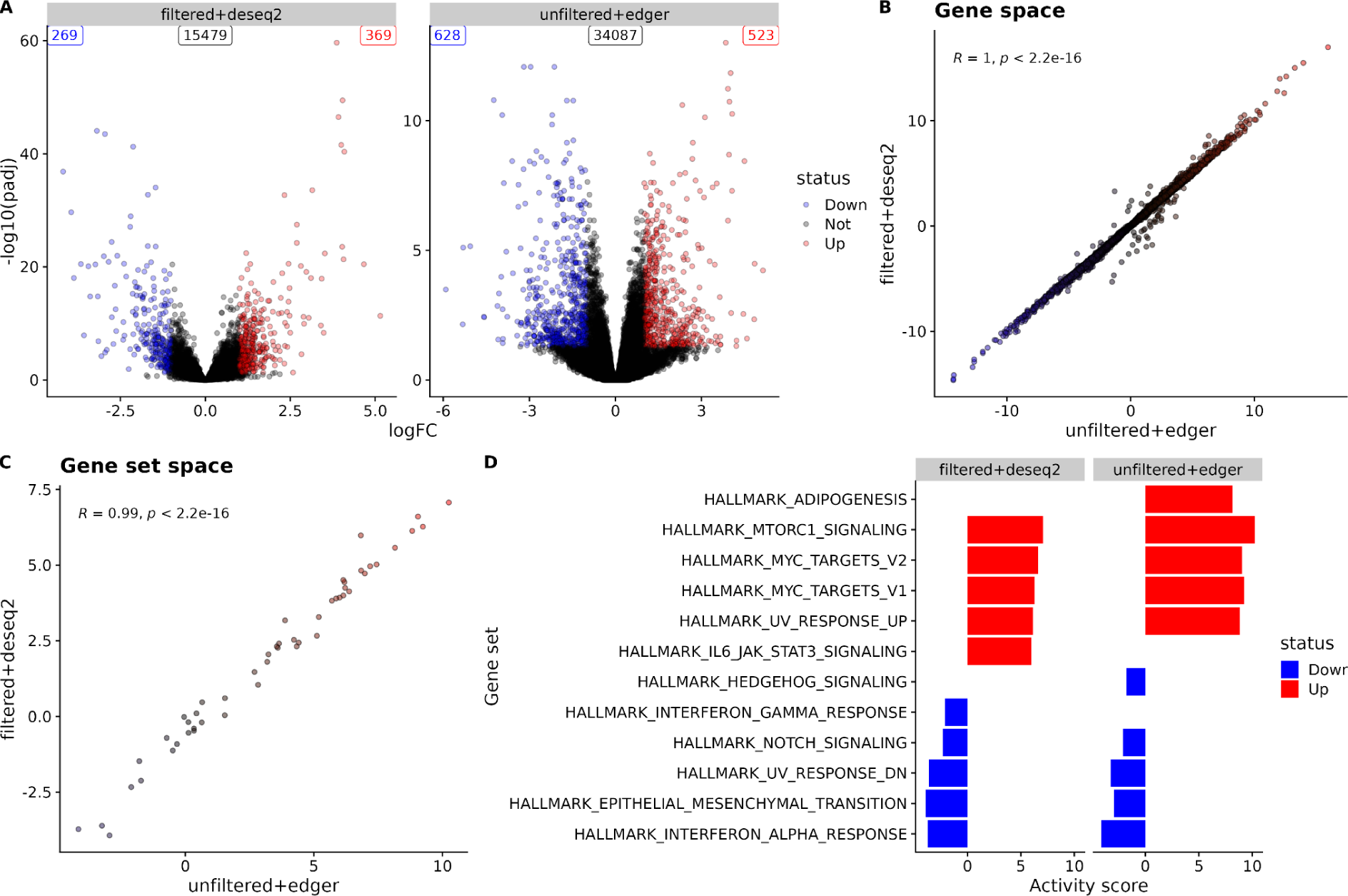
Data modalities generated and compared by FLOP. A) Volcano plots depicting the differential expression analysis results for filtered-deseq2 and unfiltered-edger in the Spurrell et al. study [44]. All genes with an adjusted P value < 0.05 and an absolute logFC > 1 are highlighted either in red (up-regulated) or blue (down-regulated). The total number of up, down and not regulated genes is displayed in the top corners. B) Scatter plot displaying Spearman correlation between the statistical estimate of both pipelines in the common gene space (each point represents a gene). The Spearman correlation score and its P value are indicated in the top left corner. C) Same scatter plot as in B, but in the gene set space (each point represents a MSigDB hallmark gene set). D) Top 10 up and down regulated gene sets per pipeline. The barplot length indicates the gene set activity score as estimated by the ulm method and the colour indicates the mode of regulation (red and blue for up-and down-regulated terms, respectively).

FLOP translates the differential expression results to the gene set space using an enrichment method available in the tool decoupleR [25]. This has two main advantages. First, differences between the pipelines are summarised in a low dimensional space that can consider information beyond the common gene space. For instance, if one pipeline filters certain genes while other doesn’t, their results are comparable in the gene set space, though not at the gene level. Second, given that FLOP focuses on gene sets, it compares pipelines on the space that is commonly used for downstream interpretation of the results. For this particular pipeline comparison, the Spearman correlation in the gene set space resembles what is observed in the common gene space (Spearman score = 0.99) (Figure 2C).

In the fifth module, FLOP isolates the top up-and down-regulated terms arising from the functional analysis, and scores its overlap.While overall agreement in the entire list of gene sets may be substantial, discrepancies can arise, particularly at the extremes of the list. To address this potential discrepancy, FLOP assesses the overlap between the extreme ends of the gene set lists across different pipelines. In this example, the Spearman correlation values for both the gene and gene set spaces exhibit high concordance. Nevertheless, when closely examining the top 10 deregulated terms, two terms differ between the two lists, resulting in a similarity score of 0.8. This indicates that 8 out of the potential 10 terms are common between the pipelines (Figure 2D). These differences can be explained by examining the statistical estimates of genes constituting these pathways (Supplementary Figure 1A). We found that genes with low expression levels significantly impact the results. These genes are ignored in the filtered+deseq2 process but are taken into account in the unfiltered+edger pipeline. Specifically, these low-expressed genes show a significant decrease in the t-value in the HALLMARK_IL6_JAK_STAT3_SIGNALING pathway (Supplementary Figure 1B). This leads to a lower score for this pathway for the unfiltered+edger pipeline. We observed similar results when we evaluated different pipelines in the same dataset (Supplementary Figure 2) and when we employed a different dataset (Supplementary Figure 3).

The aforementioned example describes one dataset, two pipelines, and one gene set collection. When we applied FLOP to the studies from the ReHeaT resource, we evaluated 5 different datasets, 12 alternative pipelines and 3 different gene set collections. This resulted in a total of 234 pipeline comparisons, each of them evaluated with the module 4 (Spearman correlation) and module 5 (top/bottom features overlap). We summarised the results of module 4 across studies for every pipeline comparison in the gene space (DE), transcription factor space (dorothea), hallmark space (msigdb_hallmarks) and signalling pathway space (progeny) (Figure 3).

**Figure 3.**
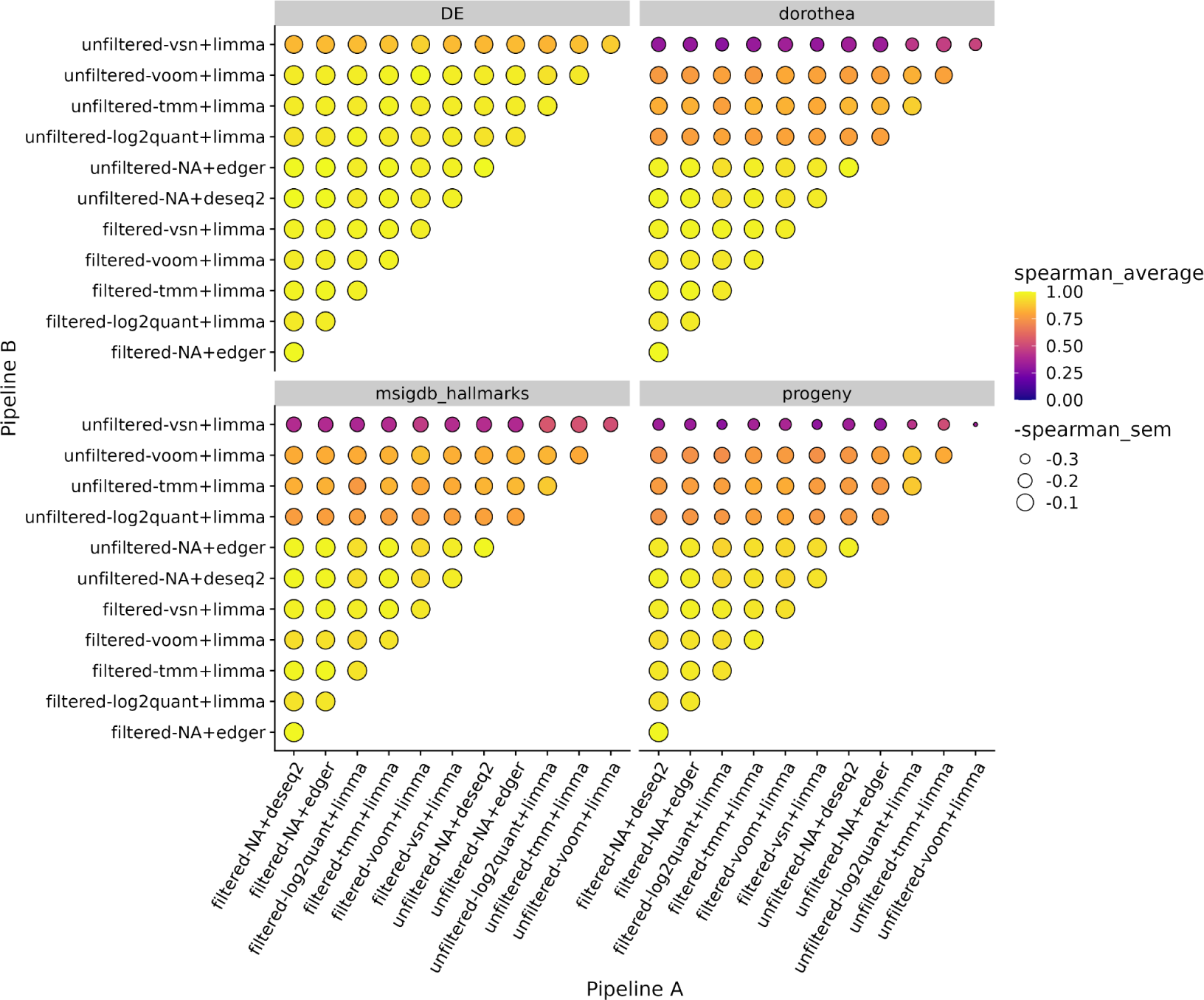
Module 4 results for ReHeaT [40] studies. Results are presented as a dotplot, with colour representing average correlation between two pipelines across studies, and with point size being inversely proportional to score standard deviation (a larger point size indicating lower variance of the correlation scores across studies). The plotting facets separate the different investigated spaces: DE for differential expression analysis results, dorothea for transcription factors, msigdb_hallmarks for transcriptional signatures and progeny for signalling pathways.

Our observations revealed that pipelines employing unfiltered counts and limma showed lower Spearman correlation with other pipelines in the gene set spaces compared to the DE space (Wilcoxon one-tailed p-value of 2.2e-16, comparing correlation values between the DE space and the gene set space). This decline in correlation values was less pronounced for unfiltered pipelines using edgeR or DESeq2 (Wilcoxon FDR-adjusted p-value 0.25, same comparison) or for pipelines employing filtered counts (Wilcoxon FDR-adjusted p-value 0.01, same comparison) (0.996, 0.993, 0.992 and 0.976 in the DE space, dorothea, msigdb_hallmarks and progeny, respectively, for unfiltered comparisons using DESeq2 and edgeR; 0.978, 0.974, 0.963 and 0.956 in the DE space, dorothea, msigdb_hallmarks and progeny, respectively, for filtered comparisons). Furthermore, we identified that the pipeline using unfiltered counts, VSN for normalisation, and limma for DE analysis exhibited notably lower Spearman correlation values with all other pipelines, both in the DE and functional spaces.

It’s important to highlight that if these differences were solely due to the smaller observation count for gene-set space correlation calculations, such an effect would be consistent across all pipeline comparisons, which is not what we observe. These findings demonstrate the utility of FLOP in quantifying the potential variability resulting from the selection of different pipelines on the agreement among the complete lists of gene sets.

We then evaluated the output of module 5 for the same cohort of studies (Figure 4). Overall, we found that similarity scores are less stable, and display lower values than Spearman correlation scores across studies (Supplementary Figure 4). In other words, focusing on the top up and down regulated gene sets, which is common practice in the interpretation of functional results, revealed a lower level of agreement between pipelines than when the entire lists of gene sets were considered. At the pipeline level, we found similar trends to those observed in the module 4 output, that is, pipelines that employ unfiltered counts and limma for DE have an overall lower Spearman correlation compared to the gene set space (Wilcoxon FDR-adjusted p-value 0.3618), and the pipeline ‘unfiltered+vsn+limma’ shows the worst similarity scores compared to all the other pipelines (0.481, 0.364, 0.447 and 0.591 in the DE space, dorothea, msigdb_hallmarks and progeny, respectively, for comparisons involving unfiltered+vsn+limma; 0.631, 0.595, 0.664, 0.734, in the DE space, dorothea, msigdb_hallmarks and progeny, respectively, for all unfiltered comparisons involving limma; 0.719, 0.712, 0.785, 0.825, in the DE space, dorothea, msigdb_hallmarks and progeny, respectively, for the rest of comparisons). These results highlight the critical role of selecting appropriate preprocessing methods to ensure more consistent and reliable downstream functional interpretations, particularly when focusing on the most deregulated gene sets.

**Figure 4.**
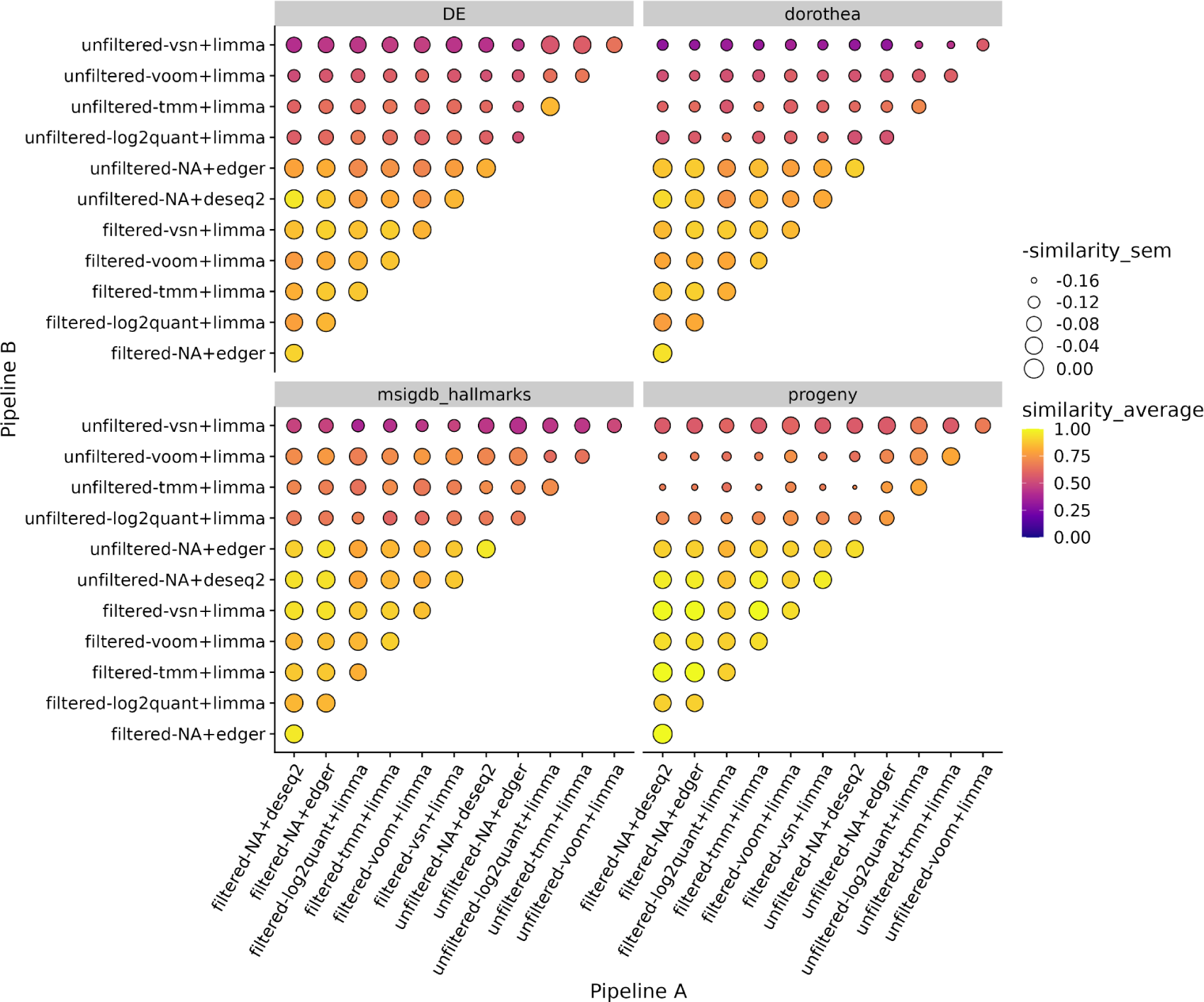
Module 5 results for ReHeaT [40] studies. Results are presented as a dotplot, with colour representing average similarity between two pipelines across studies, and with point size being inversely proportional to its standard deviation. The plotting facets separate the different investigated spaces: DE for differential expression analysis results, dorothea for transcription factors, msigdb_hallmarks for transcriptional signatures and progeny for signalling pathways.

### 2.3 Application of FLOP to perturbed and basal transcriptomic profiles of cancer cell lines

Following the application of FLOP to ReHeaT studies, we sought to demonstrate its utility in additional biological contexts. To do so, we selected two resources that profiled cancer cell lines from different angles. First, we applied FLOP to data from the Cancer Cell Line Encyclopedia (CCLE) [41]. Briefly, we retrieved the basal transcriptomic profiles for 1019 cell lines, grouped them in 18 tissues of origin, and carried out all potential non-redundant tissue comparisons. This resulted in a total of 153 contrasts, with 20 cell lines per tissue (Supplementary Table 2). Next, we retrieved transcriptomic data from the recently published PANACEA DREAM challenge [42]. In this study, a total of 11 cancer cell lines were treated with 32 different kinase inhibitors (Supplementary Table 3), with the aim of evaluating approaches to predict drug mechanisms of action from perturbational transcriptomic data. We performed all the potential contrasts between treated and untreated samples for every cell line. This resulted in 352 contrasts, with an average number of samples of 47 and 2 for untreated and treated samples, respectively (Supplementary Table 2).

After applying FLOP, we conducted an analysis of the output from modules 4 and 5. It is important to note that while in ReHeaT, each Spearman correlation data point between pipelines represented an independent study, in PANACEA and CCLE, each data point corresponding to one contrast within a specific dataset. In the context of module IV, we observed varying results depending on the dataset being evaluated. For CCLE, we found that most Spearman correlation values were well conserved, as supported by the data. However, in PANACEA, we observed a pattern similar to what was previously observed in ReHeaT, where pipelines utilising unfiltered counts displayed lower and less stable correlation values (Figure 5).

**Figure 5.**
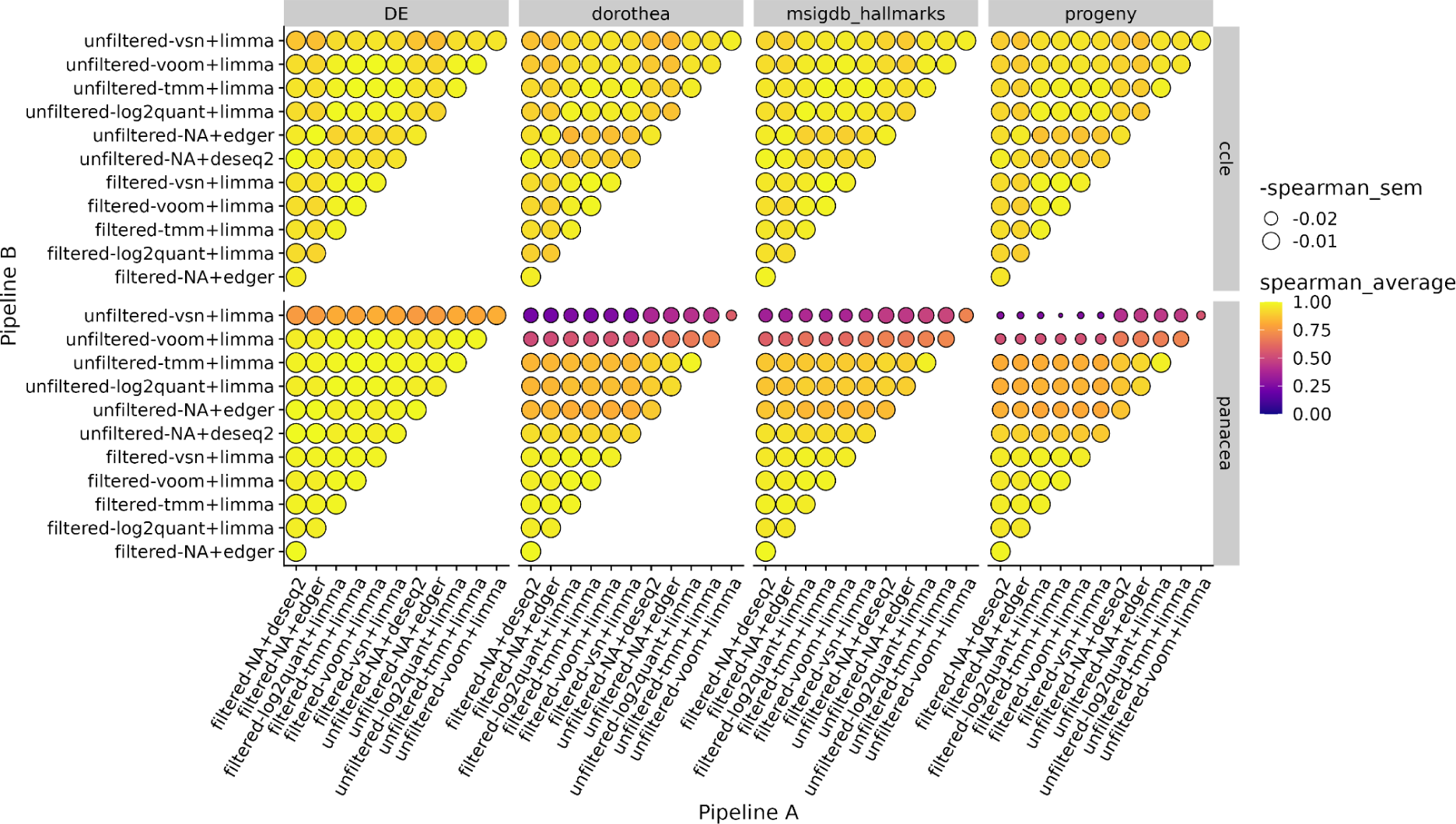
Module 4 results for CCLE [41] and PANACEA [42] studies. Results are presented as a dotplot, with colour representing average Spearman correlation between two pipelines across contrasts, and with point size being inversely proportional to its standard deviation. The plotting facets separate the investigated spaces vertically and the different datasets horizontally.

We also observed a similar trend in the output of module V, although similarity values were in general much lower, even between filtered pipelines, (Supplementary Figure 5). We attribute this difference to the varying number of samples per group in each dataset and the type of contrasts being conducted, which may result in higher statistical power in CCLE compared to PANACEA.

Overall, these findings emphasise the potential impact of selecting specific preprocessing methods on downstream functional results, which may not be apparent when only considering the differential expression (DE) space. It further highlights the utility of FLOP in scenarios with limited statistical power, enabling researchers to make more informed decisions regarding pipeline selection and functional analysis.

## 3. Discussion

In this study, we present FLOP, a workflow designed to assess the impact of different transcriptomic data analysis pipelines on downstream enrichment analysis results.

We applied FLOP to five studies sourced from ReHeaT, a meta-resource of heart failure transcriptomics data developed by our group, which represents a common clinical application scenario of transcriptomic data analysis. We showcased its capabilities to detect instances of low agreement between pipelines in the gene set space, which might not be present in the differential expression analysis space. Through the analysis of FLOP’s module 5 output, we found that, while agreements between entire gene set lists are generally high, they can be notably lower when focusing on the extremely ranked features.

We also applied FLOP to cancer cell line data from two distinct studies, namely, CCLE [41] and the PANACEA DREAM challenge [42]. These datasets were chosen for their anticipated differences in statistical power due to variations in biological contrasts explored and sample size. As expected, the results obtained in the CCLE dataset exhibited higher robustness across different pipeline choices. Conversely, the results derived from the PANACEA dataset displayed much greater variability, both when considering the entire gene set list and when focusing on its extremes.

Numerous benchmarks have investigated the impact of method choice on differential expression (DE) analysis results [14,15,26–33]. However, none of these studies evaluated the impact of method selection on downstream gene set enrichment results. Additionally, various pipelines have been developed to handle both DE analysis and functional analysis tasks. For instance, in [48], authors present a pipeline for pathway enrichment analysis on RNA-seq data. However, it lacks the flexibility to choose from different preprocessing methods. Similarly, the tool described in [49] primarily focuses on network analysis of RNA-seq data but also offers the capability to perform GO overrepresentation analysis. Yet, it provides only limited options in terms of method combinations. In contrast, [50] proposed a pipeline that allows for differential expression analysis using several methods, but it lacks a comprehensive start-to-end workflow to handle the entire process and does not offer a means to effectively compare the methods based on their results.

In this context, FLOP stands out as a unique and valuable tool that addresses all these limitations. It offers researchers the ability to assess the impact of different pipelines on downstream enrichment analysis results, thus enabling a more comprehensive understanding of the implications of method selection. In addition, it is essential to recognize that most transcriptomic studies are often conducted in settings with limited statistical power. The results in the PANACEA dataset illustrate the value of FLOP to guide pipeline selection, assess its impact, and prevent the pursuit of potentially misleading enrichment analysis results. By using FLOP to evaluate pipeline performance in datasets with varying statistical power, researchers can gain a more comprehensive understanding of the reliability and consistency of functional analyses. This empowers researchers to make informed decisions in scenarios with limited statistical power, ultimately enhancing the credibility of enrichment analysis and facilitating more accurate biological insights from transcriptomic studies.

This study has some limitations: First, FLOP per se cannot determine which pipeline performs better; it is designed to quantify differences between pipelines. We considered using simulated data, but we faced challenges in creating accurate count-level data simulations of the downstream effects for hallmarks, TFs, or signalling pathways. We also examined predefining gene sets for specific datasets, but we were concerned about the generalizability of the conclusions drawn that way. Thus, we opted to remain unbiased, focusing on quantifying inconsistencies among pipelines instead of attempting to rank them. Second, for our applications, we arbitrarily defined certain parameters that can influence the comparison outcomes. Some examples include the minimum number of counts per sample required for the filtering module or the number of top deregulated terms considered when evaluating similarities. Third, due to the combinatorial complexity involved, we focused primarily on preprocessing and employed a single method for enrichment analysis (ulm). However, the assessment of other enrichment methods could be easily implemented through DecoupleR. While it may come with higher computational costs, it could provide further valuable insights into the interactions of normalisation, DE, and functional analysis methods.

In its initial release, FLOP specifically targets bulk RNA-Seq data analysis. We plan to expand its scope to include single-cell RNA-Seq and proteomics data in a subsequent iteration. Given the rapidly evolving landscape of single-cell data analysis, it may take some time before it stabilises enough to support the use of a method like FLOP. For instance, benchmarks evaluating alternative normalisation methods for single-cell data are still under active development [35,51].

In summary, we believe that FLOP makes a valuable contribution to the transcriptomics data analysis community. First, we have packed many alternative preprocessing pipelines in a single one-liner workflow, easy to configure and run. Very often, methods are selected in regard to the lab’s expertise, previous experience, availability of documentation or ease of installation. FLOP overcomes all these challenges and provides an easy-access interface to many state-of-the-art pipelines. Second, we expect FLOP to foster the reproducibility of functional results across the community, or at least to increase the awareness of the lack of it. FLOP’s metrics are context-independent and easy to interpret, enabling users to quantify the impact of method choices in the results that they report. Thus, FLOP paves the way for a more robust and reproducible functional analysis of transcriptomics data.

## 4. Methods

### 4.1 Installation, code availability and reproducibility

FLOP can be installed from GitHub (<u>Link</u>) or Zenodo (<u>Link</u>). We packed all the required dependencies in a conda environment which needs to be set up before running the workflow. The exact list of requirements can be viewed in this yaml file (<u>Link</u>).

A test dataset can be found in Zenodo (<u>Link</u>). It is a small subset of PANACEA containing three treatments and one cell line. The seven datasets used to perform the analysis detailed in this manuscript were also deposited in Zenodo (<u>Link</u>), along with the files containing the prior knowledge sources as to 31.08.2023 (<u>Link</u>). A one-liner option is available to perform an end-to-end analysis using the datasets and prior knowledge sources deposited in Zenodo and obtain the figures present in this manuscript.

### 4.2 Data preprocessing

We applied FLOP to 8 different datasets (Supplementary Table 2): 5 datasets from the ReHeat resource, the PANACEA DREAM challenge and CCLE. We used 5 datasets from the ReHeaT resource [40]: Spurrell19 [44], LiuR [43], Pepin19 [45], Schiano17 [46] and Yang14 [47]. These datasets contain information about patients who suffered heart failure, along with other metadata such as age, gender, etc.

We downloaded the PANACEA gene counts from the NCBI GEO portal (GEO accession number: GSE186341) [42]. The dataset contains samples from 32 treatments in 11 cell lines. We performed contrasts between each treatment (2 samples) and DMSO (28-60 samples) for each of the 11 cell lines, resulting in 352 comparisons (Supplementary Table 3). We translated the gene ID’s from ENTREZ IDs to gene symbols. In multi-mapping situations where several ENTREZ IDs were pointing to the same gene symbol, counts were summed.

The CCLE dataset contains RNA-seq data from 1019 cancer cell lines, which are grouped by 26 parental tissue types [41]. We obtained the raw gene counts from the DepMap portal (<u>Link</u>). We removed tissues with less than 20 cell lines (8 tissues). To account for large differences in cell line number by tissue of origin, we randomly subsetted 20 cell lines per each of the 18 remaining tissues. We then performed all non-redundant pairwise contrasts between all tissue types, resulting in 153 comparisons.

### 4.3 Workflow

The count matrix contains genes as rows and samples as columns, and the metadata file specifies the experimental groups to which each sample belongs and the covariates to consider during the filtering and differential expression analysis. Users can also provide a contrast list, which specifies which sample groups to compare. If a contrast list is not provided, all possible non-redundant pairwise comparisons are performed by default.

FLOP integrates several methods for the normalisation, differential expression (DE) and functional analysis of RNA-seq data in a modular way. To increase computational efficiency, scalability and reproducibility, we embedded the methods in a Nextflow workflow. The workflow is divided into several processes. The first process downloads the three gene set collections: PROGENy [35], DoRothEA [36] and MSigDB hallmarks [37]. The second step carries out the 6 alternative DE analysis pipelines: tmm-limma, vsn-limma, voom-limma, log quantile-limma, edgeR and DESeq2. The next process merges the 6 pipelines’ output files into a long-format table, which serves as input for the next process, decoupleR. After the functional analysis, several consecutive modules merge the results of each dataset. Lastly, the two evaluation modules perform the analysis detailed in following sections: Rank correlation analysis and Top/bottom features overlap analysis.

FLOP output consists of four files: the differential expression results, separated by pipeline and contrast, the activity scores and p-values for all gene sets, and the results for modules 4 and 5. We implemented two profiles to control the parameters for the Nextflow run, depending if the workflow is going to be run on a desktop computer or in a slurm-controlled HPC environment.

### 4.4 Filtering strategies

To filter lowly expressed genes, we used the function *filterByExpr*, from the edgeR package, which eliminates genes that are not sufficiently expressed in a minimum number of samples per group. In other words, it takes into account the experimental design and differentiates between genes lowly expressed across groups (probably not biologically relevant) and genes lowly expressed in only some groups but not in others (potentially relevant) [10]. We used the default parameters implemented in this function for all the datasets included in this study (Supplementary Table 4).

To prevent artificial noise in the activity scores of the functional terms, we filtered out contrasts which did not have a sufficient signal, understood as a relevant number of DE genes per contrast, by implementing a cutoff of a minimum of 30 DE genes (this value can be customised).

### 4.5 Normalisation and DE analysis

We selected four methods for the normalisation of raw counts (vsn, TMM, log quantile normalisation and voom), and three methods for DE analysis (limma, DESeq2 and edgeR). We applied limma on the normalised values, while we applied DESeq2 and edgeR directly on non-normalized counts, since these methods perform an internal normalisation strategy prior to the differential analysis instead. edgeR did not provide t-statistics, unlike limma or DESeq2 (we treated Wald statistics as t-like statistics). Therefore, we transformed edgeR F statistics into t-statistics via the relation t^2^ = F for a valid comparison between methods. Parameter settings and package versions can be found in the Supplementary Table 4.

### 4.6 Functional analysis

decoupleR [25] implements multiple strategies to carry out the functional analysis of omics data by combining it with prior knowledge in the form of gene sets. Here, we used t-values as input for the analysis. Using the python version of decoupleR, we applied the univariate linear model (ulm) method. We used the default threshold of a minimum of 5 genes per gene set.

We used three prior-knowledge resources: PROGENy [37], DoRothEA [38] and MSigDB hallmarks [39]. PROGENy is a PK resource that provides a collection of responsive genes for 14 different pathways by analysis of a large set of signalling perturbation experiments. The MSigDB Hallmarks are a collection of 50 independent gene sets that encompass genes which represent a well-defined biological state or process. DoRothEA is a resource that provides relationships between 1541 human transcription factors and their target genes. We downloaded all the gene sets via decoupleR (Supplementary Table 4).

### 4.7 Evaluation modules

In module 4, we apply Spearman rank correlation to the gene set enrichment results generated by each pair of pipelines and prior-knowledge source.

In module 5, we take the top N up-regulated and N down-regulated functional categories to calculate the similarity index: for DoRothEA, N = 15; for MSigDB hallmarks, N = 5; and for PROGENy, N = 3. N was defined according to each gene set collection size. For genes, the top 5% and bottom 5% of the total number of genes were used as N. The similarity index is calculated using the formula 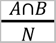, being A∩B the intersection of two lists of elements, and N the total number of elements in the lists. A similarity score of one indicates that both pipelines have the same top and bottom terms in their ranked lists, whereas a score of zero indicates no overlap between these sections of the ranked lists.

### 4.8 Statistical analysis

All p-values detailed in the results section were obtained via a one-tailed Wilcoxon rank sum test. We provide FDR corrected p-values in both the text and in FLOP results [52].

## 5 Acknowledgements

MGR was supported through state funds approved by the State Parliament of Baden-Württemberg for the Innovation Campus Health + Life Science Alliance Heidelberg Mannheim. RORF is supported by DFG through CRC/SFB 1550 “Molecular Circuits of Heart Disease”. We thank Jan Lanzer for providing us with count-level ReHeaT RNA-Seq data and metadata.

## 7. Authors contributions

**VP:** Data Curation, Formal Analysis, Investigation, Methodology, Resources, Software, Writing – Original Draft Preparation.

**AG:** Formal Analysis, Methodology, Validation, Writing – Original Draft Preparation.

**PBM:** Resources, Methodology.

**RORF:** Conceptualization, Investigation, Resources, Project Administration.

**JT:** Formal Analysis, Validation.

**MGR:** Conceptualization, Formal Analysis, Investigation, Methodology, Visualization, Supervision, Project Administration.

**JSR:** Supervision, Project Administration, Funding Acquisition.

## Supplementary Materials

### Supplementary Figures

**Supplementary Figure 1:**
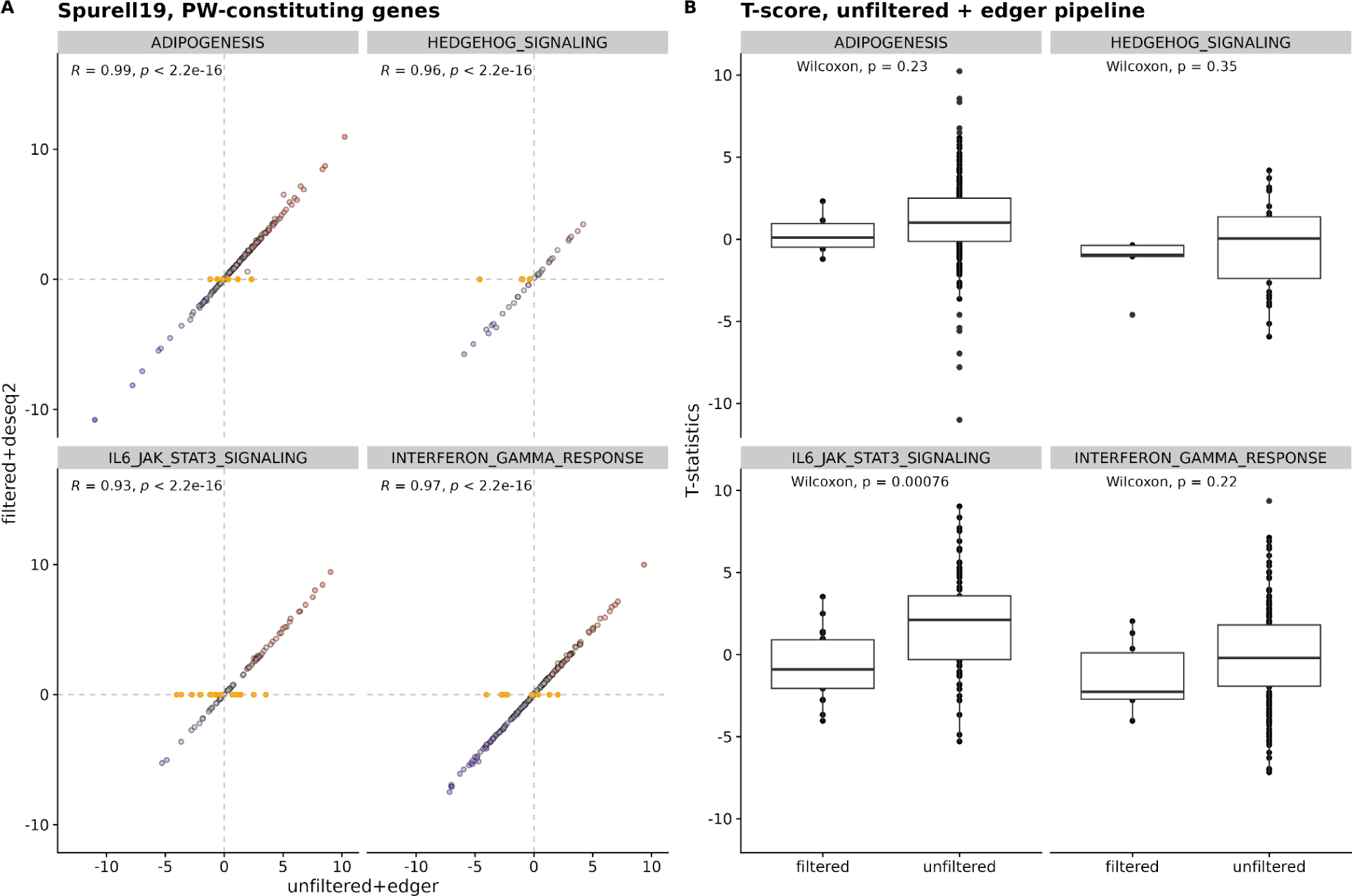
Impact of lowly expressed genes on gene set enrichment results. A) differential expression statistics (t-values) are compared for the two pipelines across the genes constituting the genesets. B) Comparison of filtered and unfiltered pathway constituting genes. Filtered genes are associated with lower t-statistics than unfiltered genes, thus they decrease the geneset scores for the pipeline where they are considered (unfiltered + edger).

**Supplementary Figure 2:**
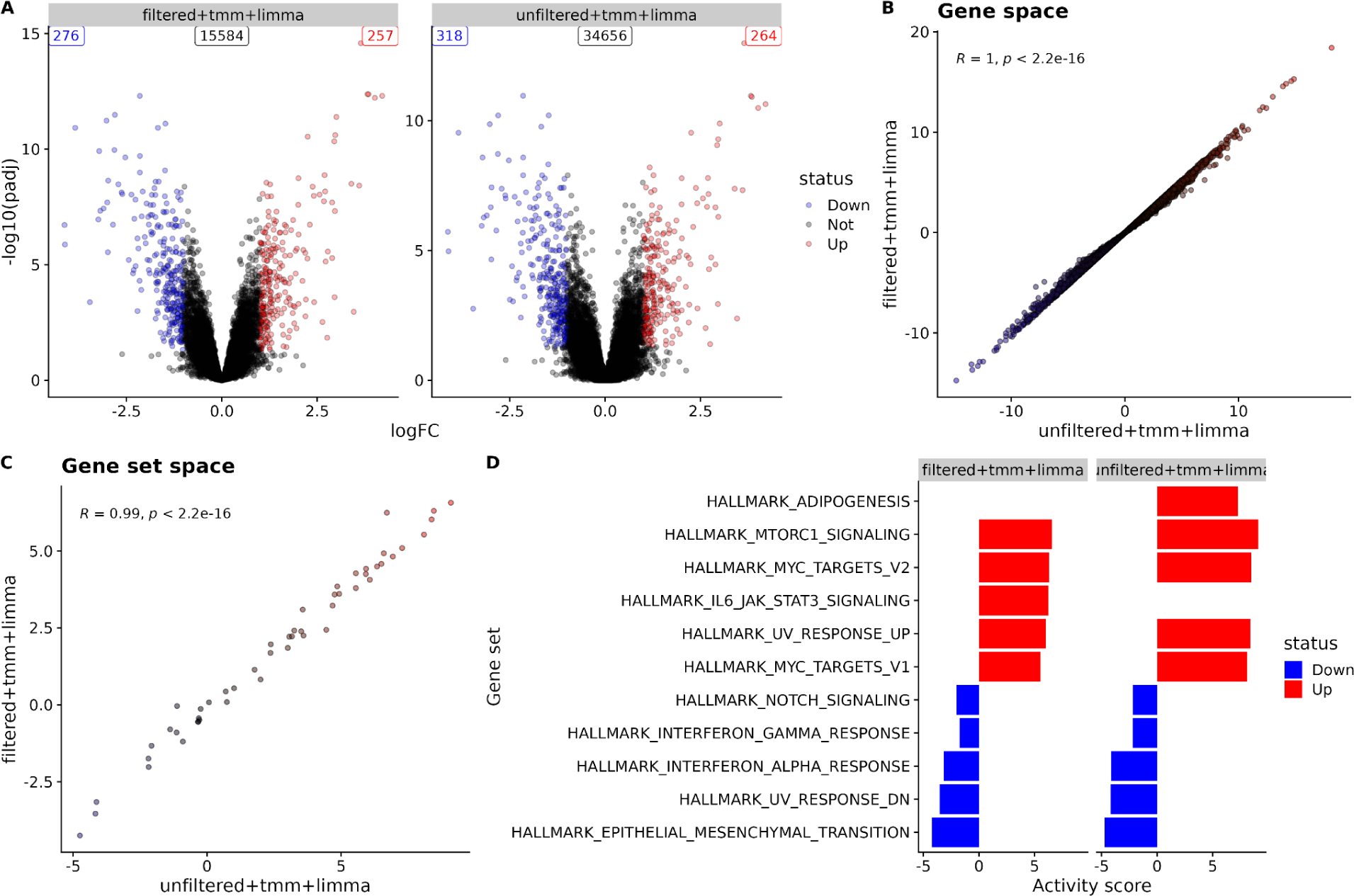
Data modalities generated and compared by FLOP. A) Volcano plots depicting the differential expression analysis results for filtered-tmm-limma and unfiltered-tmm-limma in the Spurrell et al. study [44]. All genes with an adjusted P value < 0.05 and an absolute logFC > 1 are highlighted either in red (up-regulated) or blue (down-regulated). The total number of up, down and not regulated genes is displayed in the top corners. B) Scatter plot displaying Spearman correlation between the statistical estimate of both pipelines in the common gene space (each point represents a gene). The Spearman correlation score and its P value are indicated in the top left corner. C) Same scatter plot as in B, but in the gene set space (each point represents a MSigDB hallmark gene set). D) Top 10 up and down regulated gene sets per pipeline. The barplot length indicates the gene set activity score as estimated by the ulm method and the colour indicates whether the mode of regulation (red and blue for up- and down-regulated terms, respectively).

**Supplementary Figure 3:**
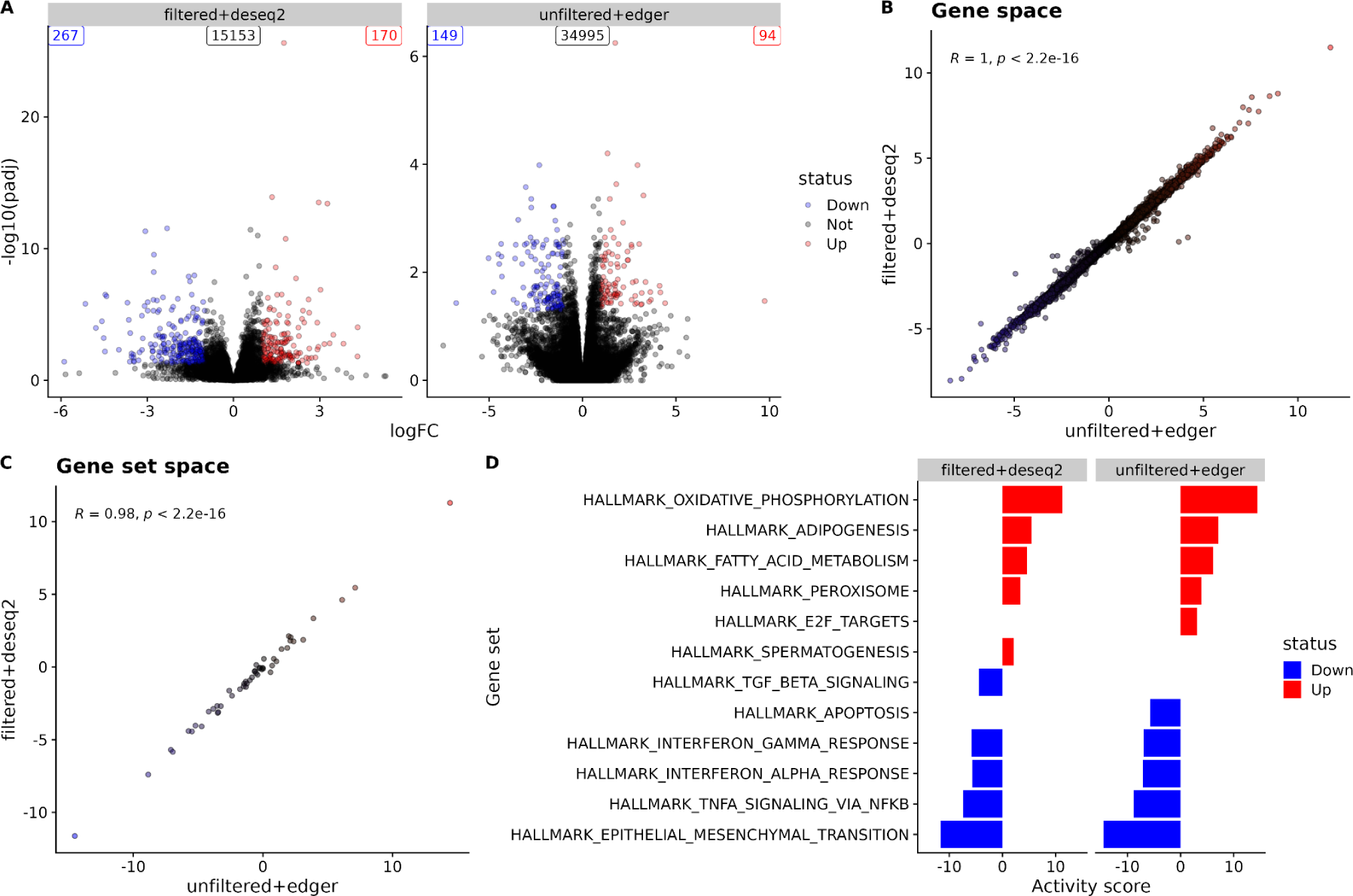
Data modalities generated and compared by FLOP. A) Volcano plots depicting the differential expression analysis results for filtered-deseq2 and unfiltered-edger in the Yang et al. study [47]. All genes with an adjusted P value < 0.05 and an absolute logFC > 1 are highlighted either in red (up-regulated) or blue (down-regulated). The total number of up, down and not regulated genes is displayed in the top corners. B) Scatter plot displaying Spearman correlation between the statistical estimate of both pipelines in the common gene space (each point represents a gene). The Spearman correlation score and its P value are indicated in the top left corner. C) Same scatter plot as in B, but in the gene set space (each point represents a MSigDB hallmark gene set). D) Top 10 up and down regulated gene sets per pipeline. The barplot length indicates the gene set activity score as estimated by the ulm method and the colour indicates whether the mode of regulation (red and blue for up- and down-regulated terms, respectively).

**Supplementary Figure 4:**
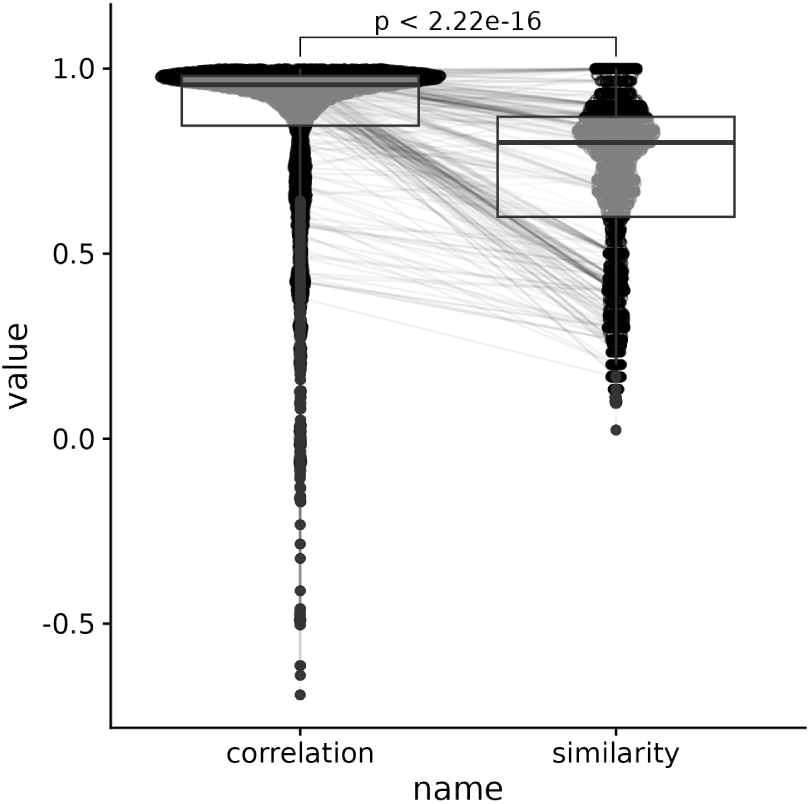
boxplot showing Spearman rank correlation values and similarity values between pipelines for all 5 ReHeaT datasets [40]. Overall, we found that similarity scores are less stable, and display lower values than Spearman correlation scores across studies

**Supplementary Figure 5:**
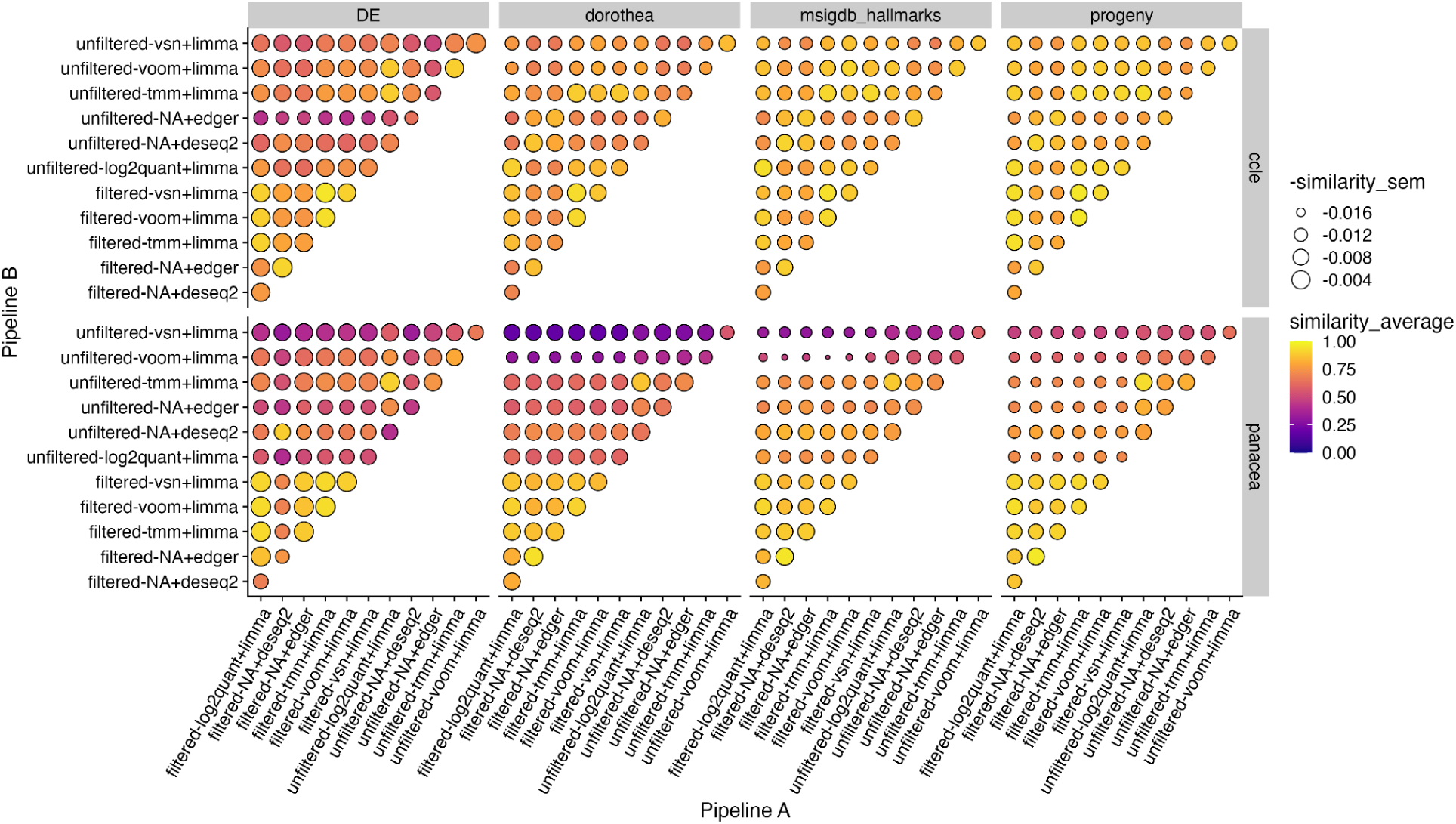
Module 5 results for CCLE [41] and PANACEA [42] studies. Results are presented as a dotplot, with colour representing average Spearman correlation between two pipelines across contrasts, and with point size being inversely proportional to its standard deviation. The plotting facets separate the investigated spaces vertically and the different datasets horizontally.

### Supplementary Tables

**Supplementary Table 1.** <u>List of benchmarking studies.</u> The table contains the name, the DOI of the published paper, a short summary of the methodology used and the types of data used in the analysis.

**Supplementary Table 2:** Overview of the datasets used in this study: number of total samples per contrast, separated by control and treatment (if applicable), number of comparisons and the covariates used in the differential expression and the filtering modules. DCM refers to disease cardiomyopathy, HTx refers to if the biopsy was collected from explanted hearts or after introducing an assisting device.

| Dataset | N samples<br>per contrast | Ctrl samples<br>per contrast | Treatment/<br>Disease samples<br>per contrast | N contrasts | Covariates |
| --- | --- | --- | --- | --- | --- |
| Spurrell19 [44] | 33 | 18 | 15 | 1 | - |
| Liu R [43] | 5 | 3 | 2 | 1 | Age, gender |
| Pepin19 [45] | 9 | 3 | 6 | 1 | Age, gender, DCM |
| Schiano17 [46] | 6 | 4 | 2 | 1 | - |
| Yang14 [47] | 24 | 8 | 16 | 1 | Age, gender, DCM |
| PANACEA [42] | 30-62 | 28-60, avg 47 | 2 | 352 | - |
| CCLE [41] | 40 | 20 cell lines per tissue |  | 153 | - |

**Supplementary Table 3.**
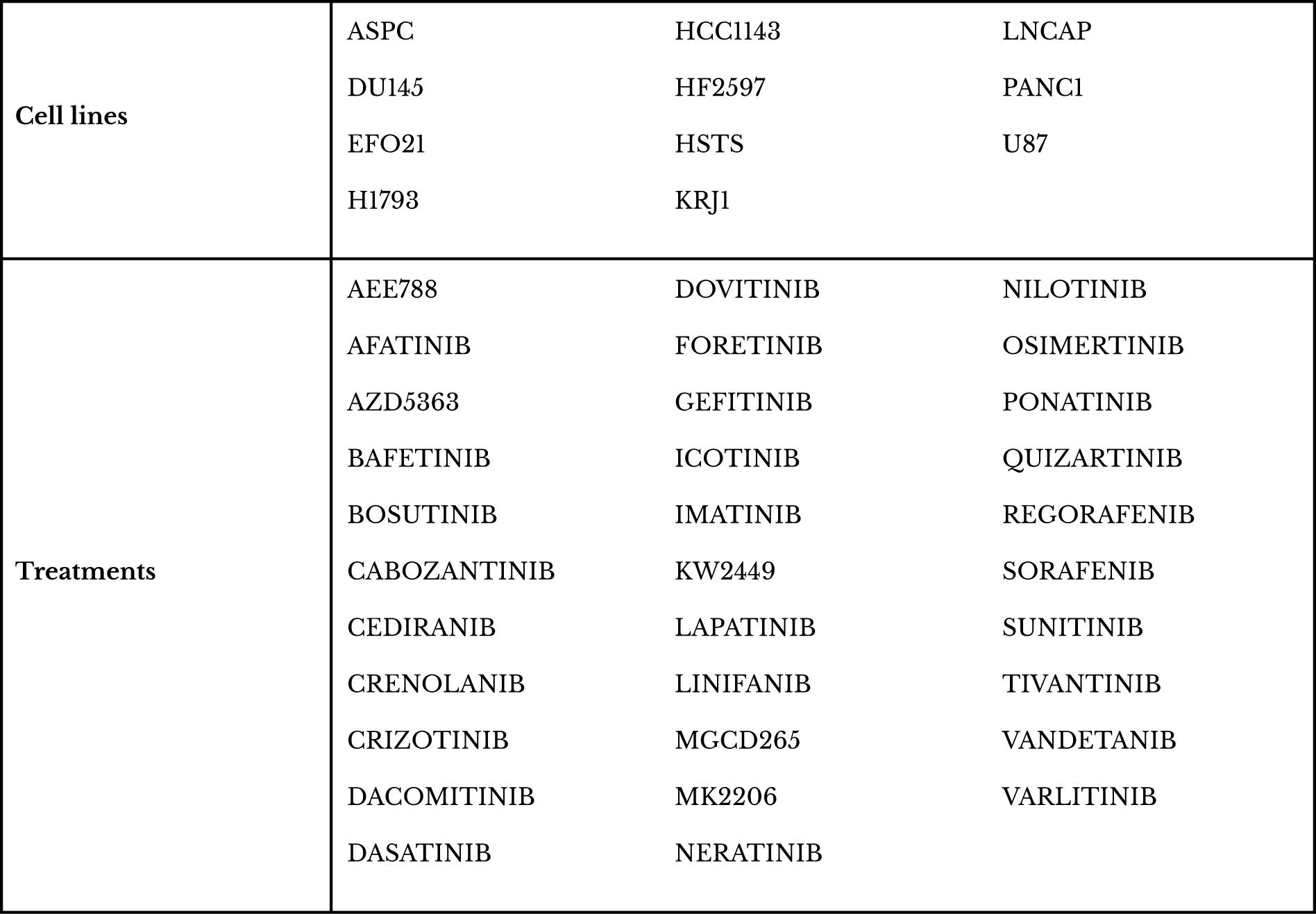
List of cell lines and treatments in the PANACEA dataset. Contrasts were produced comparing each treatment against DMSO (control), for every cell line.

**Supplementary Table 4.**
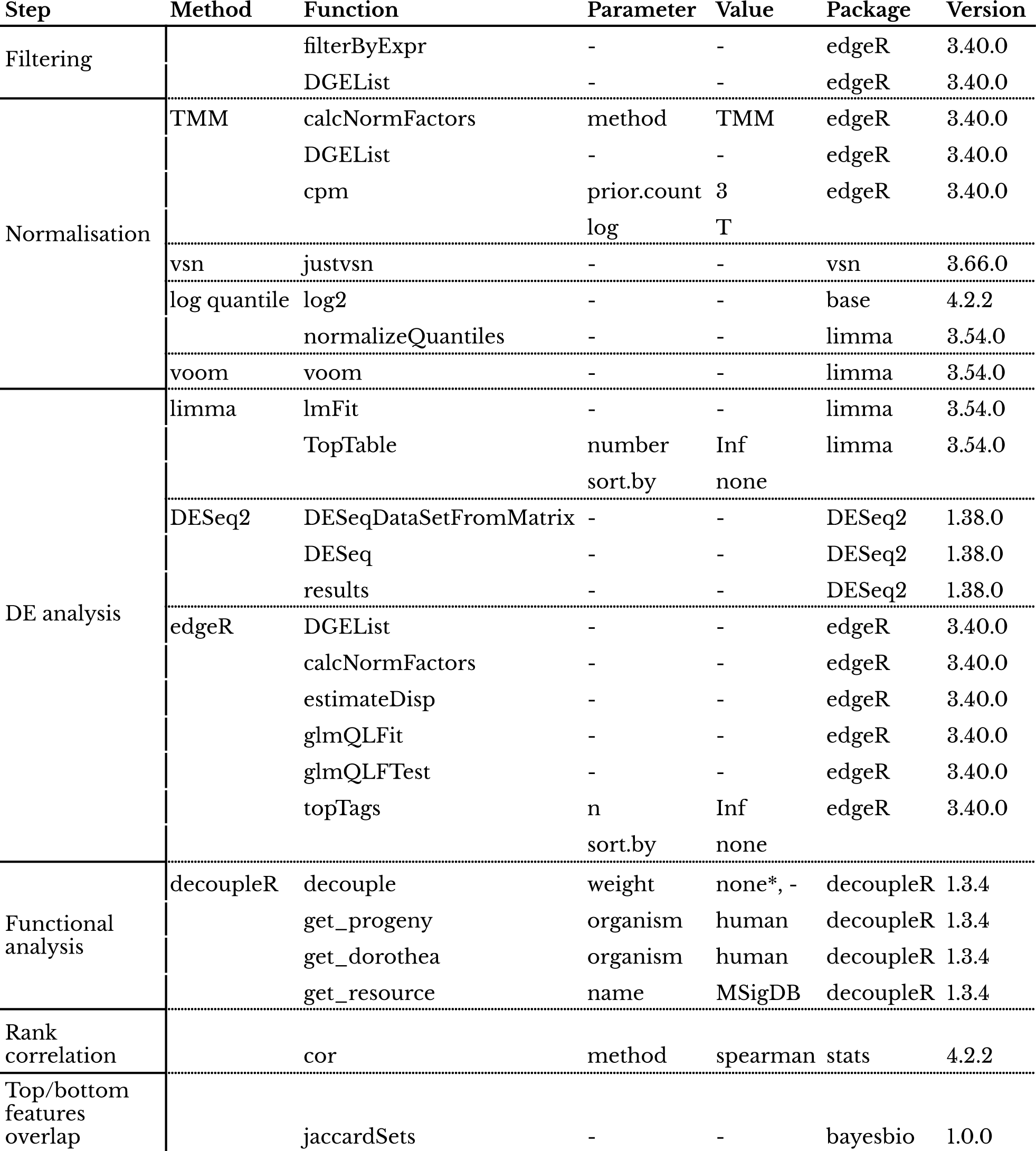
List of the most relevant functions used in FLOP. Here, we included only the parameters whose values we changed. Parameters which are not specified in the table took default values. We also included the package name and version to which the function belongs. * indicates that weights were only used in those PKs that included them.

