## Supplementary Table 1 for "Assessing the impact of transcriptomics data analysis pipelines on downstream functional enrichment results"

| Evaluated step | Study | DOI | Methods | Datasets | Evaluation strategy |
|---|---|---|---|---|---|
| Normalization | [14] | https://doi.org/10.1186/s12859-015-0778-7 | Eight non-abundance (RC, UQ, Med, TMM, DESeq, Q, RPKM, and ERPKM) and two abundance estimation normalization methods (RSEM and Sailfish) | Real Illumina high-throughput RNA-Seq of 35- and 76-nucleotide sequences produced in the MAQC project and simulation reads. | Spearman rank correlation of genes between the normalization results from RNA-Seq and MAQC qRT-PCR values for 996 genes. |
| normalization + DE | [15] | https://doi.org/10.1186/gb-2013-14-9-r95 | DESeq, edgeR, limma-QN, limma-voom, PoissonSeq, CuffDiff, baySeq | SEQC study (GSE49712) and ENCODE project | Normalization: they performed hierarchical clustering of samples (if clusters resemble biological differences, the normalization was successful) and then estimated Dunn cluster validity index.<br>DE: ROC analysis using genes previously measured by qRT-PCR as set of true differentiation (cutoff of 0.5 logfc). Also they performed an evaluation of type1 errors via null models (using technical replicate samples). For genes uniquely expressed in one condition, they assessed type II errors by performing an isotonic regression model between signal-to-noise ratio and adjusted p-value. |
| DE | [26] | https://doi.org/10.1186/s12859-018-2261-8 | ALDEX2 family of methods, DESeq2, EdgeR | two simulated RNA-seq datasets and two real datasets (SRP082682, PRJNA277985 ) | Precision and recall from a contingency table of the simulated state of differential expression (as a binary) compared with the predicted state of differential expression (as a binary). For one real dataset, they computed this per microarray channel and then did the average of both estimates. |
| DE | [27] | http://dx.doi.org/10.1038/s41467-017-00050-4 | Regarding differential expression: DESeq2, limma, edgeR, Cuffdiff, ballgown, sleuth. | 15 Illumina and Pacific Biosciences (PacBio) data sets from normal human sample NA1287810, human MCF-7 breast cancer cell, H1 human embryonic stem cell (hESC), and the Sequencing Quality Control Consortium (SEQC) data set. | Compared the detected differentially expressed genes with ground truth from expression changes measured by qRT-PCR: spearman rank correlation and RMSD between log2FC of qRT-PCR vs RNA-seq, AUC-30 scores meaning area under ROC for a false discovery rate of 30 percent. |
| DE | [28] | https://doi.org/10.1093/nar/gkv806 | ROTS, DESeq, DESeq2, edgeR, CuffDiff, limma, BaySeq, NOISeq, PoissonSeq | Spike-in dataset (GSE49712), cancer genome atlas (ccRCC), ccRCC validation dataset (EGAS00001000509) | They used predetermined fold changes to assess the sensitivity and specificity of the methods via ROC/AUC analysis and FDR. They also assessed the usefulness of ROTS to associate certain marker genes with patient outcome via a risk score. |
| DE | [29] | https://doi.org/10.1371/journal.pone.0232271 | edgeR, edgeR.glm, edgeR.rb, edgeR.ql, edgeR.ql.rb, DESeq, DESeq2, voom-tmm, voom.qn, voom.sw, baySeq, bayseq.qn, PoissonSeq, SAMseq, ROTS | SEQC study (GSE49712) as spike-in data, simulated datasets from TGCA kidney renal clear cell carcinoma/normal dataset and inbred mouse dataset (bottomly) | ROC curve, true positive rate and FDR per method, for several simulated datasets containing several combinations of dispersion, percentage of DE genes and presence or absence of weak effect sizes |
| DE | [30] | https://doi.org/10.1186/1471-2105-14-91 | DESeq, edgeR NBPseq, TSPM, baySeq, EBSeq, NOSeq, SAMseq, Shrinkseq, limma-voom, limma-vst | several simulated datasets using a negbinomial distribution, with mean and dispersion estimates from real RNA-seq data. They also tested 2, 5 and 10 samples per condition. Finally, they also used two real datasets (10.1371/journal.pone.0017820) | AUC, false discovery curves for different number of DE genes, bot in one and two directions, and for 5 samples per condition. They also checked type I error control, for pval<0.05, and FDR control, for FDR<0.05. For the real datasets, they compared the number of found DE genes per method and their overlap (venn). No ground truth was available for benchmarking here |
| DE | [31] | https://doi.org/10.1261/rna.046011.114 | DESeq, DESeq2, edgeR, EBSeq, sSeq | Six RNA-seq datasets: Bottomly, Bullard, Huang, Montgomery-pickrell, tuch, Qian, were used to estimate biological paramenters (GLM) to later generate simulated data | TPR, Matthews correlation coefficient and F-measure for several sample sizes and for 100 random samplings, |
| single-cell normalization + DE | [32] | https://doi.org/10.1186/s13059-020-02136-7 | Several methods, starting from a Seurat pipeline, in the following steps: doublet detection, filtering, normalization, feature selection, denoising, dimensionality reduction, and clustering | Two real datasets (GSE79636 via IPSCpower package, data from seqc package) and a simulated one. | Adjusted Rand index, mutual information, silhouette width, precision, recall, AUROC, running time, rate of misclassification (for doublets) |
| DE | [33] | https://doi.org/10.1371/journal.pone.0190152 | Regarding differentia expression: baySeq, DESeq, DESeq2, EBSeq, edgeR, limma-voom, NOIseq, SAMseq | Real Illumina Microarray Quality Control dataset and qrt-PCR as ground truth. | They compared identified DE genes between methods and qRT-PCR ground truth. They counted the true positives and false positives per method. They also computed ROC curve, specificity and FPR from several number of methods |
